# Biobehavioral synchrony across species: Evidence for multi-level regulatory dynamics in human–canine dyads

**DOI:** 10.64898/2026.06.24.734140

**Authors:** Miiamaaria V. Kujala, Aija Koskela, Iina Valkeajärvi, Heini Törnqvist, Virpi-Liisa Kykyri, Takefumi Kikusui, Jan Kujala

## Abstract

Coordinated dynamics between individuals are a hallmark of social interaction, yet the temporal structure and physiological basis of such coupling beyond human species remain poorly understood. Here, we investigated cross-species biobehavioral synchrony by simultaneously quantifying motion dynamics and autonomic activity with hyperscanning of human–canine dyads. We observed both spontaneous and task-related synchrony across motion dynamics, heart rate, and heart rate variability at multiple timescales. Importantly, synchrony was modulated by individual and relational factors. Task-related autonomic synchrony was affected by the human temperament, whereas greater familiarity within the dyad altered the leader–follower dynamics, shifting directional influence from human-led toward canine-driven coordination. Motion synchrony emerged with minimal delay, whereas cardiac synchrony unfolded across longer timescales, suggesting coordinated processes underlying the shared activity, arousal, and autonomic regulation. Our findings extend current models of social synchrony beyond human interactions and reveal that regulatory dynamics underlying coordinated behavior operate across species boundaries.

## Introduction

Social synchronization, a form of group cohesion present in spatial, hormonal and neuronal mechanisms, exists across animal species (1). In humans, such alignment of behavior and physiology is seen across diverse contexts, from rhythmic joint actions to the coordination of goal-directed movements and spontaneous transference of affective states (2). Although early work has focused on structured tasks, recent evidence has revealed the emergence of spontaneous interpersonal synchrony via natural movement (3). These observations imply that synchrony reflects a general principle of social interaction that may extend beyond conspecific relationships (4). However, the regulatory dynamics of distinct biobehavioral mechanisms within such cross-species interactions remain poorly understood.

Recent data show hormonal and behavioral correlations across species, including human-canine dyads that share a long coevolution (5, 6). Human traits further link with canine behavior (7, 8). Neuroticism, or negative affectivity, drive the correlated hair cortisol levels within dyads (6), whereas extraversion relates to better dog trainability (9). However, these studies are based largely on observational approaches, lacking temporally resolved measures of interaction dynamics used to study human interpersonal synchrony (10). How the precise temporal patterns of cross-species dyadic synchrony unfold under different conditions remains unknown, along with the direction of actions. Furthermore, existing studies typically examine single domains, leaving the multiple biobehavioral levels of movement, arousal, and autonomic regulation unresolved. Thus, a unified, multi-level account of dynamic cross-species coordination is lacking.

Here, we investigated the emergence and temporal alignment of cross-species biobehavioral synchrony with motion dynamics and autonomic nervous system hyperscanning. We examined three explanatory levels operating at fundamentally different temporal scales: motion dynamics (MOV), heart rate (HR), and heart rate variability (HRV), quantifying both spontaneous and task-related synchrony, scrutinizing the leader-follower dynamics and examining the influence of behavioral and psychological variables on cross-species synchrony.

## Results

### Clustering and spontaneous synchrony

The synchrony effect sizes across the five conditions (rest and four tasks) and three measures (MOV, HR, HRV), formed three data clusters in the 2D representation (**Fig. 1A**). Separate measures formed a main body of the three clusters, with mixed HR samples in the HRV cluster, HRV samples in the MOV cluster, and MOV samples from separate tasks in the HR cluster. Spontaneous synchrony of MOV, HR and HRV occurred in humans and dogs within lags up to 0, 5 and 10 seconds (*i.e*., −*L*_max_ ≤ *L* ≤ *L*_max_, where L_max_ varies across data types) for MOV, HR and HRV respectively (MOV 0s: t(28)=15.39, *p*<0.001, Cohen’s *d*=2.85, CI [2.02, 3.68]; HR 5s: t(28)=9.6, *p*<0.001, Cohen’s *d*=1.78, CI [1.19, 2.37]; HRV 10s: t(28)=3.49, *p*=0.002, Cohen’s *d*=0.65, CI [0.24, 1.04]; **Fig. 1B**). Spontaneous synchrony was statistically highly significant especially for the motion dynamics despite the low activity levels (humans: 0.016 ± 0.0003 dg/dt, dogs: 0.027 ± 0.001 dg/dt).

**Fig. 1.**
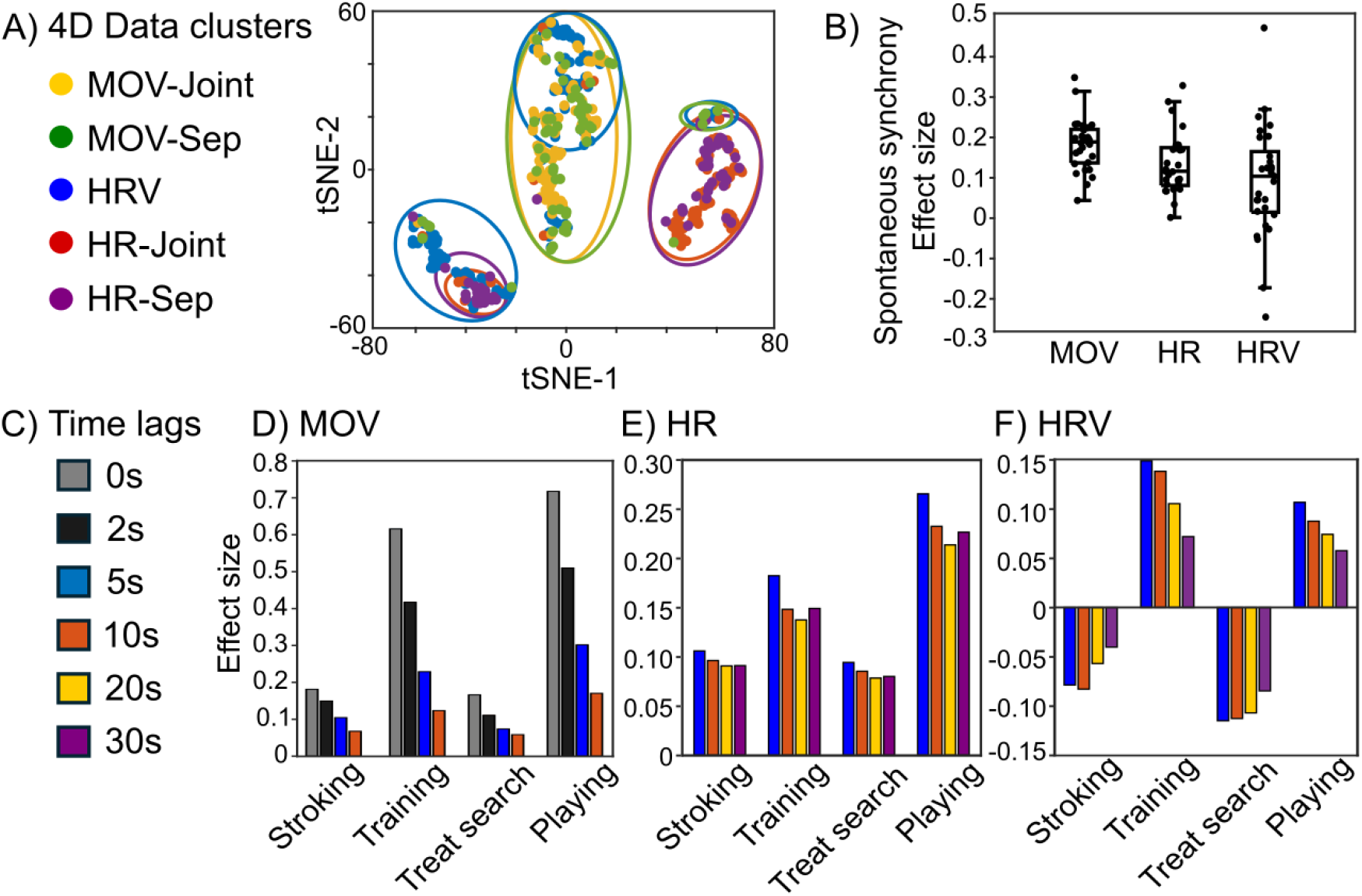
Data clusters and effect sizes in spontaneous and task-related synchrony. A) 2D tSNE representation of 4D synchrony data (four time lags) across dyads, conditions, tasks and measures. tSNE-1 = first tSNE dimension, tSNE-2 = second tSNE dimension. MOV = motion dynamics, HR = heart rate, HRV = heart rate variability, Joint = conditions with similar tasks for human and dog (rest, training, playing), Sep = conditions with separate tasks (stroking, treat search). B) Spontaneous synchrony for motion dynamics, heart rate and heart rate variability. C) Representation of SUSY for different time lags between human and canine data. D) Effect sizes of motion dynamics for the different time lags (0–10s); E) heart rate (5–30s); and F) heart rate variability (5–30s).

### Task-related synchrony

Significant positive synchrony of human–canine dyads occurred across all tasks within both motion dynamics (t(28)=6.79, *p*<0.001, Cohen’s *d*=1.26, CI [0.77, 1.75]; t(28)=8.68, *p*<0.001, Cohen’s *d*=1.61, CI [1.05, 2.16]; t(28)=5.49, *p*<0.001, Cohen’s *d*=1.02, CI [0.56, 1.47]; t(28)=10.08, *p*<0.001, Cohen’s *d*=1.87, CI [1.26, 2.47]) and HR (t(28)=2.70, *p*=0.011, Cohen’s *d*=0.50, CI [0.11, 0.88]; t(28)=3.76, *p*<0.001, Cohen’s *d*=0.70, CI [0.29, 1.10]; t(28)=2.71, *p*=0.011, Cohen’s *d*=0.50, CI [0.11, 0.89] and t(21)=3.45, *p*=0.002, Cohen’s *d*=0.74, CI [0.26, 1.20] for stroking, training, treat search and playing, respectively; **Fig. 1C-1E**). Furthermore, HRV synchrony was detected during training and treat search (training: within-phase, (t(27)=2.13, *p*=0.043, Cohen’s *d*=0.40, CI [0.13, 0.78]; treat search: anti-phase, (t(27)=-2.11, *p*=0.044, Cohen’s *d*=-0.39, CI [-0.01, -0.77], **Fig. 1F**).

### Effect of individual factors

The association between task-related synchrony and individual reactivity was examined by correlating human temperament (11) with the synchrony effect size. Human negative affectivity correlated negatively with the effect size of HR and HRV, but not motion dynamics, during the treat search task (MOV 0s: *ρ*=.13, *p*=0.512, CI [-.31, .49]; HR 5s: *ρ*=-.48, *p*=0.009, CI [-.11, -.76]; HRV 10s: *ρ*=-.42, *p*=0.024, CI [-.31, .49]; **Fig. 2A**). Thus, higher human negative affectivity led to a higher magnitude of negative synchrony in the dyadic cardiac measures. Instead, higher human extraversion correlated with the effect size of motion dynamics, but not with cardiac measures, during training (MOV 0s: *ρ*=.45, *p*=0.016, CI [.05, .73]; HR 5s: *ρ*=.15, *p*=0.460, CI [-.23, .46]; HRV 10s: *ρ*=.04, *p*=0.853, CI [-.41, .46]; **Fig. 2B**).

**Fig. 2.**
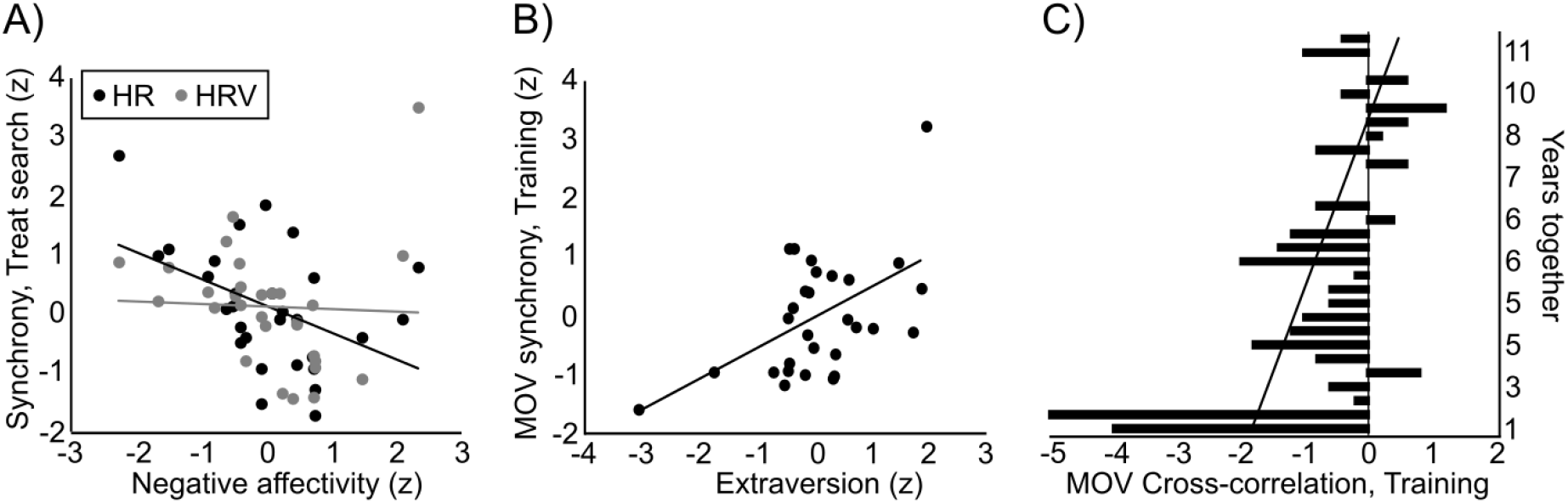
Human-dog synchrony connections with human temperament and the familiarity of the dyad. A) Caretaker negative affectivity correlates negatively with HR and HRV synchrony during Treat search. B) Caretaker extraversion correlates with MOV synchrony during Training. C) Correlation between familiarity (years together, y-axis) and the time-lag (seconds) of MOV leader-follower analysis during Training (x-axis: negative values = human leads the motion; positive values = dog leads the motion).

### Leader-follower dynamics

Training as a joint activity was scrutinized further with a cross-correlation analysis. While the cross-correlation analysis revealed a peak in human-dog synchrony generally at zero seconds, variation in the peak timing was explained by the higher familiarity of the human– canine dyad (*ρ*=.34, *p*=0.032, CI [.04, .69]; **Fig. 2C**). Thus, the higher familiarity of the dyad switched the direction of the leader of the motion dynamics from the human toward the dog, with more familiar human–canine dyads having more instantaneous motion synchrony, and in some dyads, dog leading the motion.

## Discussion

Our results demonstrate cross-species biobehavioral synchrony across multiple levels of organization. As demonstrated for human interactions (3), also familiar human–canine dyads exhibit spontaneous in addition to task-related synchronization. Task-related synchrony was consistently observed for motion and heart rate, whereas HRV synchrony was restricted to conditions involving cooperation or asymmetrical engagement. In these conditions, synchrony likely incorporates affective and regulatory processes beyond shared activity. Notably, HRV synchrony was pronounced despite the intrinsically non-linear cardiac dynamics characteristic of dogs, underscoring the robustness of cross-species autonomic coupling. Furthermore, autonomic synchrony was modulated by individual factors: during treat search, higher human negative affectivity was associated with stronger negative synchrony. This pattern may reflect complementary regulatory states, such that successful task performance by the dog reduces arousal in the human observer, particularly in individuals prone to heightened negative affect.

We also show that more extraverted humans exhibited higher motion synchrony with their dogs, and greater familiarity influenced the timing of cross-correlation during training. As extraversion is associated with positive affect (11), it may enhance social reinforcement during training. Furthermore, dogs with longer shared histories appeared better in predicting human actions, indicating learning of non-conspecific motion patterns. Dogs can predict the motion of moving objects (12), and this capacity likely extends to tracking biological motion. Although the role of familiarity in predicting non-conspecific motion has not been directly examined, more familiar human–canine dyads have shown stronger cardiac coupling during stress (13). Thus, these findings highlight the role of learning in emotional and motor alignment across species.

The examination of the effects of context, task, relationship quality, and individual factors on interpersonal synchrony between people has only begun, as has disentangling the underlying multimodality (14). We quantify these factors within a cross-species relationship, moving beyond anthropocentrism to capture the connection between humans and the biosphere. Our study indicates that cross-species synchrony reflects structured, multi-level regulatory dynamics affected by both context and individual differences. These results advance our understanding of social cohesion mechanisms and raise broader questions regarding the nature of social bonding, interspecies communication, and the evolution of affiliative behavior.

### Materials and Methods

Simultaneous measurement of electrocardiography and velocity change (3D accelerometers) were acquired from human–canine dyads while they shared a space of a 30m^2^ room, processed at three separate explanatory levels: MOV, HR, and HRV. By adopting the surrogate synchrony (SUSY) method (15), we calculated spontaneous synchrony occurring within human–canine dyads during periods of resting and task-related synchrony during four tasks requiring different levels of interaction: stroking, training, treat searching and playing. The resting condition was void of specific action instructions and comprised low activity but with access to multimodal contact. Data clustering was examined by subjecting the 4D data comprising SUSY effect sizes across four time lags to dimension reduction via t-distributed stochastic neighbor embedding (tSNE). Training was examined in detail with a cross-correlation analysis of MOV using 80% overlapping one-second time windows, and the correlations of synchrony effect sizes to distinct behavioral and psychological variables were calculated. The experiment had prior approval by the ethical board of the University of Jyväskylä (26.6.2020; #760/13.00.04.00/2020; amendment 05/2022). Human participants gave written informed consent before participating with their dogs. Experimentation was performed in accordance with the regulations instructed by The Finnish National Board on Research Integrity (https://tenk.fi/en) and the ARRIVE guidelines (https://arriveguidelines.org).

## Acknowledgments

We are grateful for Markku Penttonen and Viki-Veikko Elomaa for the advice with data synchronization, Tiia Silfverberg for the practical assistance and for Peppi Taalas for the measurement facilities. This study was supported by grants from the Research Council of Finland (#341092, #346430); Agria Djurförsäkring & Svenska Kennelklubben Forsknings Fond (#N2022-0007; #P2023-0001); and Emil Aaltonen Foundation (#230085; #250083K1).

